# Food for thought: Rosy-faced lovebirds (*Agapornis roseicollis*) are capable of associative symbol learning and inference-based quantity discrimination

**DOI:** 10.1101/2024.12.30.630847

**Authors:** Shengyu Wang, Kevin Ching Hei Lo, Verna Wing Ting Shiu, Christy Yuen Ching Hung, Emily Shui Kei Poon, Chris Newman, Christina D. Buesching, Simon Yung Wa Sin

## Abstract

Cognitive capacity for quantity discrimination is highly adaptive in various ecological contexts and subject to convergent evolution across diverse animal species, yet the underlying mechanism involved is not fully understood. Discrimination accuracy generally increases with the ratio between two quantities; however, this ability is expected to differ across ratio ranges. To test this we presented a novel symbol system to 28 rosy-faced lovebirds (*Agapornis roseicollis*), associating additive tally marks with symbols representing a one-to-one correspondence with different food quantities. Trained lovebirds could spontaneously infer the relative food quantities represented by other symbols. Lovebirds proved capable of (1) associating symbols (i.e., object-file symbolism); with (2) “more-less” quantity inference, by deducing food quantities based on their knowledge of this symbol-quantity association; and (3) enhancing their performance in relation to disparity ratio (conforming to Weber’s law) and absolute difference. Furthermore (4), the influence of food ratio and absolute difference varied with different ratio ranges. Within a small ratio range (≤ 3) discrimination performance improved with increments of ratio or absolute difference, whereas within a higher ratio range (> 3), the effect of these factors diminished. We conclude that rosy-faced lovebirds are capable of advanced numeracy and quantity discrimination, similar to larger parrot species.

## Introduction

The ability of animals to discriminate quantities in the contexts of resource acquisition, competition, and risk aversion is a highly adaptive facet of cognition, conserved under strong selection pressure [1,2]. Fundamental quantitative sensitivity is thus ubiquitous across taxa, exemplified among mammals (primates [3], cetaceans [4] carnivores [5,6], ungulates [7], and rodents [8]), birds [9], reptiles (lizards [10] and archosaurs [11]), amphibians [12], bony fish [13], sharks [14], and even invertebrates [15,16]. Judging quantities can be advantageous in multiple life contexts, including optimal foraging ([16,17]; and fundamental to marginal value theorem [19]), conspecific communication and cooperation [20,21], assessing numerical advantage in inter-group conflicts (both directly [22], e.g., call counting [23], and indirectly through assessing scent marking activity [24,25]), avian egg counting [9,26] and brood parasite avoidance [9,27], group affiliation [28] and distinguishing sex ratio in mating [29,30] and local resource competition [31] contexts, *inter alia*. These quantitative assessments also often interact with qualitative perceptual cues; for example, black-capped chickadees (*Poecile atricapilla*) encode information about both predator number and size in their alarm calls [21]. Furthermore, quantitative sensitivity is mediated by social and environmental [32] factors, and by whether the spontaneous or trained ability of individuals is being tested [33]. Consequently, it has proven challenging to derive a comprehensive and systematic understanding of quantity discrimination in different species and/or taxa.

Because of the diverse effects of context, species, and environments a broad range of (not mutually exclusive) theories have been proposed to explain specific forms of quantity discrimination [1]. The simplest form is subitizing, i.e., the ability to rapidly identify the quantity of items in a small group without counting them individually [34]. This is the first numerical ability developed among human infants [34] and provides a rapid and accurate judgement of quantities ≤ 4. Larger quantities (>4) must be counted, involving the precise enumeration of cardinal numerosities, a skill developed by human infants around the age of 9 months [35]. More sophisticated counting ability conforms to the object-file model that denotes each item by a symbol (file), enabling accurate small-number estimation (≤ 4) [36], as found in species as diverse as lizards [10], Clark’s nutcrackers (*Nucifraga columbiana*) [26], and rhesus monkeys (*Macaca mulatta*) [37]. Lastly, the analogue magnitude mechanism involves the attribution of an analogue representation proportional to the objects’ properties (e.g., number, size, length or surface). Such analogue representation has no limit, which allows the depiction of large numbers [38]. This has been established as a counting method used, for instance, by capuchin monkeys (*Cebus apella* [39]) and jungle crows (*Corvus macrorhynchos* [40]).

Within these mechanisms, the ability of animals to perform quantity discrimination tends to conform to Weber’s law [22], a psychological concept that describes how the perception of a difference in a stimulus depends on the original intensity of the stimulus [41]. In terms of quantitative sensitivity, the capacity to discriminate between two quantities thus typically increases as the ratio between them decreases (i.e., animals should perform better when discriminating between 2 and 8 items –a ratio of 0.25– than between 6 and 8 –a ratio of 0.75; [42]). Quantity discrimination is further affected by the magnitude (defined as the total amount across both quantities, e.g., the magnitude of 6 vs. 8 would be 14) and the disparity (i.e., the absolute difference between the two quantities; e.g., the disparity of 6 vs. 8 would be 2) of differences between contrasted quantities. Consequently, performance decreases as magnitude increases, disparity decreases, and as the ratio between contrasted quantities becomes more even [43]. Such ratio-based quantity discrimination has been found to transcend across vertebrate [44] and invertebrate species [16]. Nevertheless, research examining quantity discrimination patterns within identical magnitudes, i.e., exploring whether the absolute difference between two quantities, rather than quantity sets of varying ratios is decisive in animal choice, remains scarce.

The use of inferential reasoning as a process for making judgement calls based on available evidence allows animals to apply prior knowledge to decision-making in novel scenarios [45,46]. Specifically, inferential reasoning by exclusion involves deciding where a reward exists based on information about its absence [47], whereas transitive inference (TI) is a form of deductive reasoning that allows a relation between items that have not been explicitly compared before to be derived [48]. Thus, in a general form, TI is the ability to deduce that if Item B is related to Item C and Item C is related to Item D, then Item B must be related to Item D.

Both forms of inferential reasoning have been established in various mammal species [49], especially in primates [50,51], but are also features of the advanced cognitive abilities of certain bird taxa. Indeed, parrots (Psittacidae) and corvids (Corvidae) have been termed “feathered apes” [52,53] in the literature, due to their exceptional cognitive capacities, which are comparable to those observed in primates [54]. Transitive inference ability has been established among corvids [55,56], while inferential exclusion is well established among both corvids (e.g., Eurasian jays, *Garrulus glandarius* [57], New Caledonian crows, *Corvus moneduloides* [58], ravens, *Corvus corax* [59], carrion crows, *Corvus corone corone* [60]) and parrots (e.g., African grey parrots, *Psittacus erithacus* [61–63], Goffin cockatoos, *Cacatua goffini* [64], kea, *Nestor notabilis* [65]), where testing has shown that individuals are capable of inferring reward placement conditional upon knowing the emptiness of potential alternative sites. Furthermore, African grey parrots [66] and kea [67] are capable of probabilistic reasoning, inferring a sample based on prior knowledge about a population. Pepperberg [68] has also shown that African grey parrots can infer a cardinal value from its ordinal position on the numeral list.

Testing avian quantity discrimination in all possible combinations between one and five, has revealed that the performance of both African grey parrots [69] and the Eurasian jackdaw (*Coloeus monedula*) [70] improves with larger ratios. Jungle crows can even be trained to learn that a five-circle symbol on a cup indicates a reward inside, while a two-circle symbol does not [40]. Moreover, in five quantity combinations task (3 vs. 5, 4 vs. 5, 5 vs. 6, 5 vs. 7, 5 vs. 8), jungle crows attained higher accuracy with higher ratios [40]. It is thus reasonable to anticipate that parrots and corvids, may be able solve a “more-less” paradigm by inference from symbols representing specific food quantities; that is, associative symbol learning [71]. Nevertheless, only a limited number of Psittacidae species have been subject to comprehensive and detailed study, notably the African grey parrot. It remains largely unexplored whether other, especially smaller parrot species, exhibit comparable cognitive performance.

Rosy-faced lovebirds (*Agapornis roseicollis*) are highly social small-sized parrots that are native to arid regions in southwestern Africa [72]. Here, food resources vary substantially among seasons [73] resulting in rosy-faced lovebirds eating a highly selective diet while foraging in groups [74]. The ability to cope with complex environmental challenges, combined with sociality typically implies highly evolved cognitive abilities (“Cognitive buffer hypothesis” [75–77]). In addition, they are popular companion pets [78]. We thus used rosy-faced lovebirds as a model to test three predictions: 1) whether they are capable of associative learning, specifically object-file symbolism, by examining whether they can associate food presence-absence and food quantity with symbols; 2) whether rosy-faced lovebirds are capable of “more-less” quantity inference, by examining whether they can deduce food amounts based on prior knowledge of symbol-quantity association; 3) whether their discriminative ability improves with disparity ratio, conforming to the predictions of Weber’s law. We then explore what underlying quantity discrimination mechanism is being used by this species.

## Materials and Methods

### Study animals and housing conditions

Experiments were conducted using 28 adult rosy-faced lovebirds (19 males and 9 females, aged 2 to 3.5 years old; see Table S1) housed individually in wire-mesh cages (60cm × 40cm × 40cm) under a 12h:12h light cycle (6500K illumination; LED T5 tube, 7W, SUNSHINE) at the Centre for Comparative Medicine Research (CCMR) animal facility at the University of Hong Kong. The ambient temperature was kept at 22-24°C and humidity between 50-60%. Artificial food pellets (Mazuri Small Bird Maintenance Diet 56A6) and UV-filtered water were available *ad libitum* and changed daily.

### Experimental protocol

The same experimental setup was used to first investigate associative symbol learning, inference, and then symbol-indicated quantity discrimination ability. Each bird was tested individually in their own cage by providing a perch (length = 30 cm) with a stainless-steel cylindrical cup (diameter × height: 5cm × 5cm) attached to each end (Fig. 1a). A camera (STARCAM CB71-C / STARCAM CB73 Mini Battery IP Camera) was placed on top of the cage to record bird behaviours. Birds were able to move freely around the cage and access both cups equally from the perch. These cups were covered with tightly fitted white paper lids (2.3g/100cm^2^), upon which symbols were printed using red food-grade dye. In addition, laminated vertical signs with identical corresponding symbols were affixed behind each cup (Fig. 1a). We used a circle (i.e., ‘O’, 23mm in external diameter and 17mm in internal diameter; hereafter referred to as 0) to indicate an empty cup and different numbers of bars (i.e., ‘|’, measuring 23mm in length and 3mm in width, separated by a gap of 2 mm), to indicate (as a tally) specific food reward portions concealed under the white paper lid in each respective cup (Fig. 1b). For instance, one bar (i.e., ‘|’) indicated one sunflower seed and one food pellet in the cup, whereas 2 bars ‘||’ indicated two sunflower seeds and two food pellets in the cup etc. For clarity, we use Arabic numerals hereafter to represent all symbols in the text.

**Figure 1.**
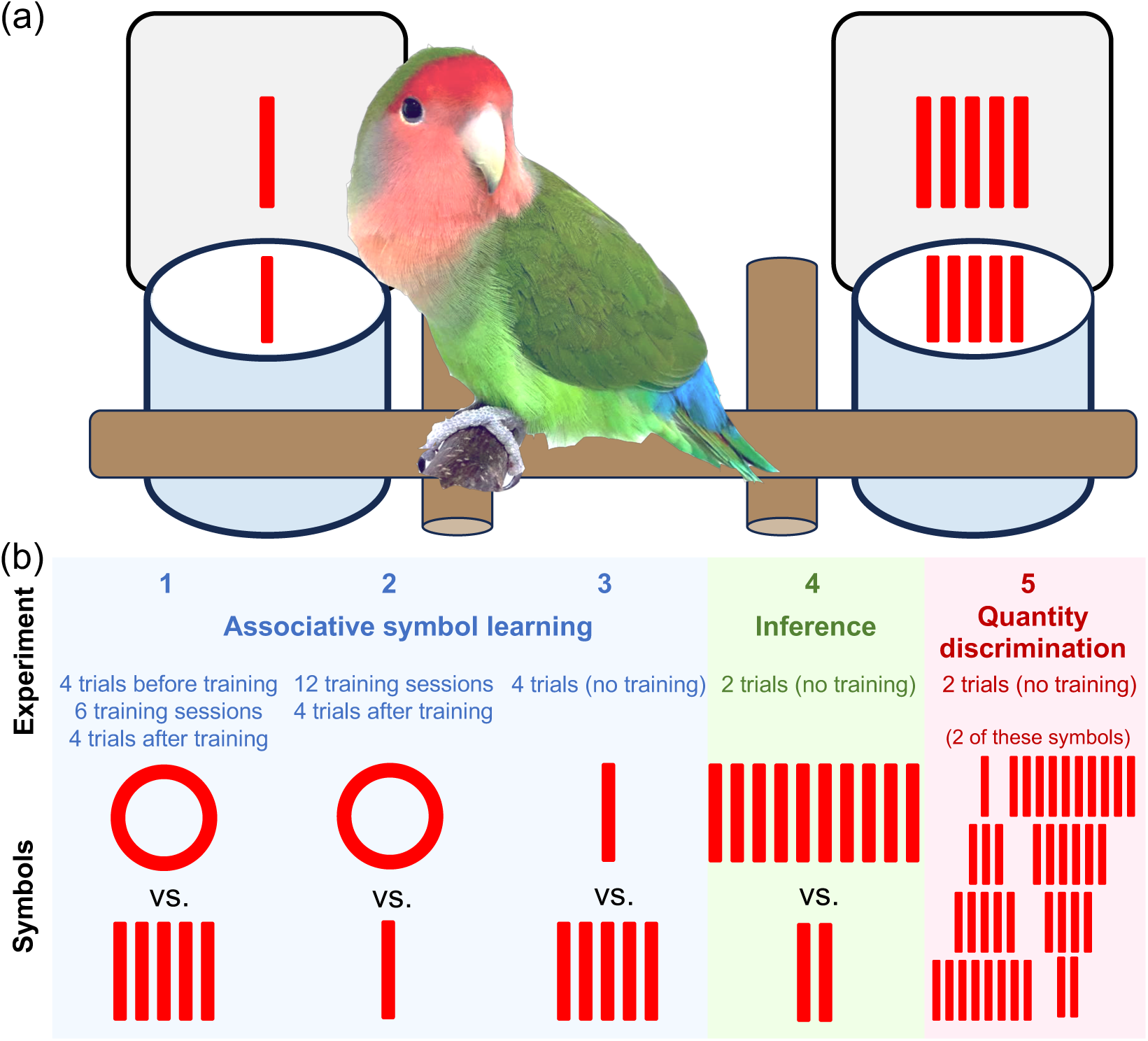
Schematic overview of the experimental design. (a) Experimental protocol. Food rewards were hidden in stainless steel cups during trials. Symbols were shown on and behind the cups. (b) The five experiments conducted on associative symbol learning, inference, and quantity discrimination, and the symbol combinations used. Experiment 1: 0 vs. 5, 4 trials before training + 6 training sessions + 4 trials after training. Experiment 2: 0 vs. 1, 12 training sessions + 4 trials after training. Experiment 3: 1 vs. 5, 4 trials (no training). Experiment 4: 2 vs. 10, 4 trials (no training). Experiment 5: 2 trials for each combination of symbols (see Table 1).

**Table 1.** Symbol combinations tested for quantity discrimination.

| Absolute<br>difference | Ratio |  |  |  |  |  |  |
| --- | --- | --- | --- | --- | --- | --- | --- |
|  | 1:1.25 | 1:2 | 1:3 | 1:4 | 1:5 | 1:8 | 1:10 |
| 9 |  |  |  |  |  |  | 1 vs. 10 |
| 8 |  |  |  |  | 2 vs. 10 |  |  |
| 7 |  |  |  |  |  | 1 vs. 8 |  |
| 6 |  |  |  | 2 vs. 8 |  |  |  |
| 5 |  |  |  |  |  |  |  |
| 4 |  | 4 vs. 8 | 2 vs. 6 |  | 1 vs. 5 |  |  |
| 3 |  |  |  | 1 vs. 4 |  |  |  |
| 2 |  |  | 1 vs. 3 |  |  |  |  |
| 1 | 4 vs. 5 | 1 vs. 2 |  |  |  |  |  |

To motivate birds to participate in the discrimination trials, we closed their feeders for 1.5 hours before the start of each trial as well as for 20 minutes after the conclusion of the experiment, as lovebirds have been shown to learn and adapt to this imposed fasting regime by eating food when available during the trial [79]. We then simultaneously attached both experimental feeder cups to the perch in the subject parrot’s cage, and gave the bird 10 minutes to tear the paper lid off the cup(s) and consume the food therein, after which both cups were removed. We defined that the bird made a choice if the bird had torn the paper off a cup and visually inspected the food contained inside. None of these birds had previously participated in any symbol-discrimination test. We then conducted a sequential series of experiments to examine the ability of lovebirds for associative symbol learning, inference, and the discrimination of quantities.

#### i) Associative object symbol learning

Before commencing these associative learning trials, we first established whether lovebirds had any intrinsic preference for a particular symbol. For this, we conducted four trials presenting cups with 0 against cups with 5, controlling for left-right side bias and potential morning-afternoon differences (i.e., the position of each cup/symbol was switched in the two trials in the morning, 11:00-12:00, and in the two trials in the afternoon, 16:00-17:00).

We then trained the birds to associate the symbol 0 with the absence of food and the symbol 5 with the presence of abundant food, to test the ability of lovebirds to form an association between specific symbols indicating the presence (in one cup) or absence (in the other cup) of food (i.e., inference by exclusion through object-file association). During this training period, we removed ca. 1/6 of the paper lid area without damaging the printed symbols so that lovebirds were able to see the content inside the food cup to inform their choice. Nevertheless, lovebirds still needed to exert effort to tear the paper lid away to fully access the food. Six rounds of training were conducted for this 0 vs. 5 symbol combination, and the symbols were provided three times in each position (i.e., left or right).

After the training sessions we conducted four experimental trials to test whether the lovebirds had learned to associate symbols with the presence or absence of food (Figure 1b, Experiment 1) by presenting 0 vs. 5 (supplementary video 1), with cups fully covered with paper lids bearing corresponding symbols. As during training sessions, the four trials were controlled for left-right side bias and potential morning-afternoon differences.

We then repeated the same training protocol as above for the symbol combination 0 vs. 1 (supplementary video 2), followed by four experimental trials for 0 vs. 1 (Figure 1b, Experiment 2; supplementary video 3). Twelve rounds of training were conducted for this 0 vs. 1 symbol combination, due to the low interest birds had in small food rewards, with the symbols provided six times in each position (i.e., left or right).

After the 0 vs. 1 trials we performed a second experiment to test whether lovebirds had learned to associate the different symbols (1 vs. 5) with different quantities of food. For this we conducted four trials of 1 vs. 5 without providing any additional training (Figure 1b, Experiment 3).

#### ii) Inference

To examine whether lovebirds could infer the meaning of additive tally mark bars comprising symbols subsequently presented to them (i.e. relative food quantities) based on the symbols they had learned during the associative learning trials, we conducted two experimental trials using combinations of two new compound symbols (i.e., 2 vs. 10), where the numbers of bars in each symbol was double that of the previously learned symbols (i.e., 1 and 5) while adhering to the same ratio disparity (Figure 1b, Experiment 4). This experiment was conducted during the morning session only to exclude performance difference between morning and afternoon sessions.

#### iii) Quantity discrimination

To investigate what mechanism lovebirds use to discriminate quantities, ratio disparities or absolute amounts, we provided food quantity combinations at disparity ratios of 1.25, 2, 3, 4, 5, 8, and 10 (Table 1), as well as two combinations with different absolute differences in each of the ratios 2, 3, 4, and 5. We ran two trials for each symbol combination, with different left-right positions to control for potential side-bias. The symbol position in each combination was randomised in the first trial. All trials were carried out in the morning. The use of the same symbol in two consecutive combinations, and any combination consisting of a familiar symbol alongside a novel additive tally bar symbol were avoided. In ca.1% of all trials, the bird did not participate (i.e., showed no interest in either feeder). In these cases we repeated the trial only once, and the trial was recorded as a missing value if the bird also did not participate in this second reiteration.

### Statistical analyses

Analyses were performed and plotted using RStudio (Version 4.2.2) [80]. We ran generalised linear mixed models (GLMM) using the R package lme4 v1.1-31 [81] to examine the influence of symbol position (left/right) on the feeder cup (left/right) chosen by birds in each combination, including bird ID and trial ID as random effects, i.e. model 1) choice position ∼ symbol position + (1|ID) + (1|trial). We then ran beta regression models using the R package betareg v3.1-4 [82] to investigate the effects of food quantity ratio and absolute difference on the proportion of trials in which a bird selected the most rewarding feeder cup as its first choice: model 2) the proportion of first choices selecting the greater food reward ∼ ratio; model 3) the proportion of first choices selecting the greater food reward ∼ ratio * ratio level (“low” when ratio ≤ 3, “high” when ratio > 3); model 4) the proportion of first choices selecting the greater food reward ∼ absolute difference; and model 5) the proportion of first choices selecting the greater food reward ∼ ratio * absolute difference.

We also tested whether an individual’s performance was influenced by symbol familiarity (number of times it had seen each symbol before testing a symbol combination) and the test order of symbol combinations: model 6) the proportion of first choices selecting the greater food reward ∼ familiarity; and model 7) the proportion of first choices selecting the greater food reward ∼ trial order). We used ANOVA to examine differences in latency to choose (time spent until the first cup was opened) across different symbol combinations: model 8) latency to choose ∼ combination. A linear model was used to investigate for any association between an individual’s performance and its sex, age, and weight: model 9) individual proportion of first choices selecting the greater food reward across all symbol combinations (morning sessions only) ∼ sex, age, and weight.

## Results

### Associative symbol learning

Prior to 0 vs. 5 cup/symbol combination training the position of symbols had no effect on the food cup chosen by lovebirds, with either symbol selected with almost equal likelihood (symbol 5 preferred over symbol 0 in 49.5% of trials; *P* = 0.694; Fig. 2; Table 2). After training, however, the proportion of first choices in which the symbol indicating the food reward cup was selected increased to 71.2% (*P* < 0.001; Fig. 2; Table 2). Similarly, after 0 vs. 1 symbol combination training lovebirds chose the cup marked with the symbol indicating a food reward significantly more often (60.6%) than they selected the empty symbol (*P* < 0.02; Table 2), although the proportion of first choices of the symbol indicating a food reward was lower than in the 0 vs. 5 trials (Table 2). In the 1 vs. 5 trials, even without any prior training lovebirds selected the symbol indicating the greater food reward significantly more often (65.2%) than the symbol indicating the cup containing less food (*P* < 0.002; Table 2).

**Figure 2.**
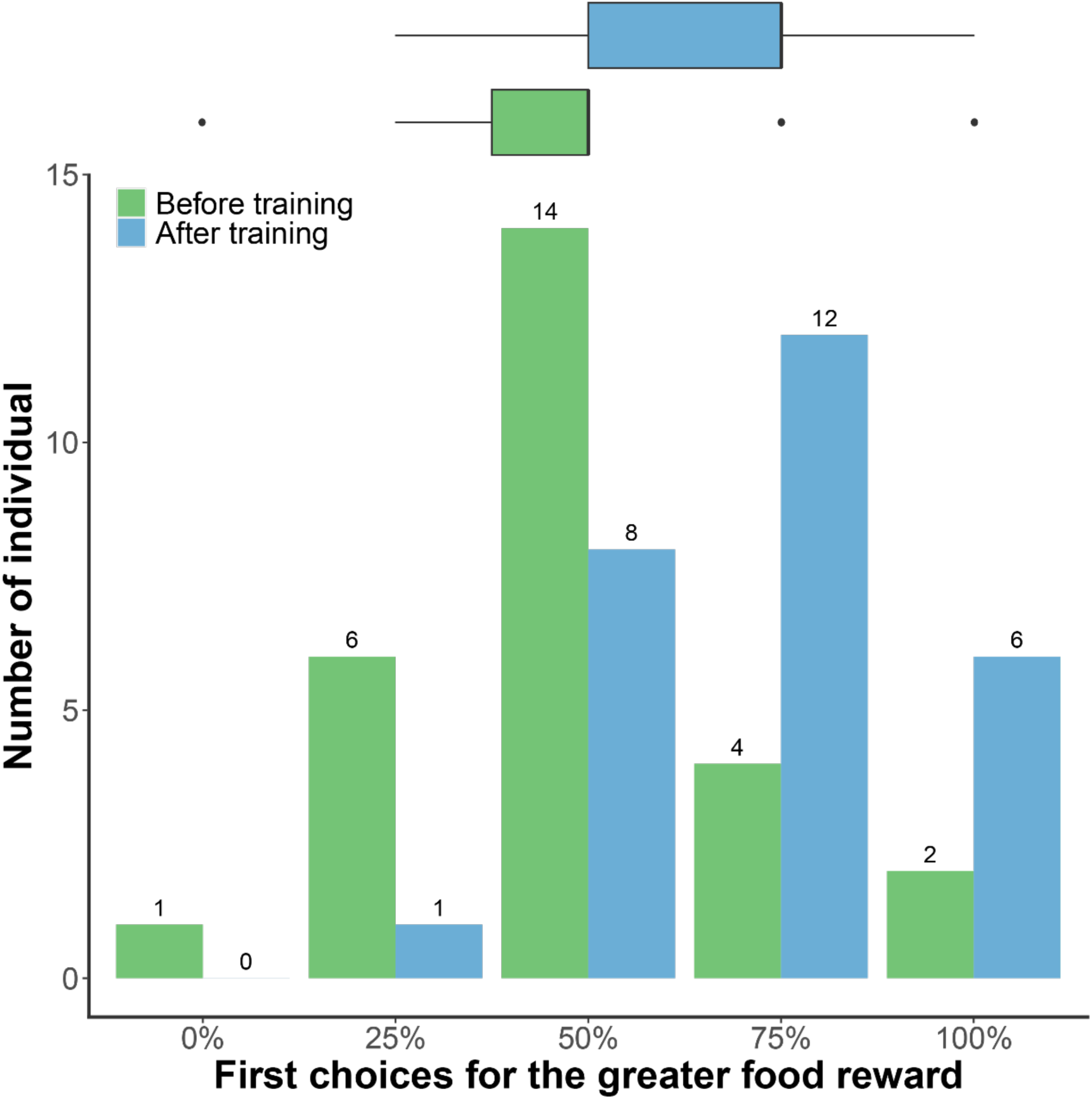
More individuals chose the cup with the greater food reward after training.

**Table 2.** Performance in associative symbol learning and inference. Asterisks indicate significant results: * *P* < 0.05; ** *P* < 0.01; *** *P* < 0.001.

| Combinations | Proportion of first choices for the greater food reward | Estimate | $P$ |
| --- | --- | --- | --- |
| 0 vs. 5 before training | 49.5% | -0.182 | 0.694 |
| 0 vs. 5 after training | 71.2% | 2.249 | <0.001*** |
| 0 vs. 1 after training | 60.6% | 1.331 | 0.012* |
| 1 vs. 5 | 65.2% | 1.537 | 0.001** |
| 2 vs. 10 | 67.9% | 1.708 | 0.019* |

### Inference

When lovebirds were presented with the new symbol combination of 2 vs. 10, they were significantly more likely to choose the symbol indicating the greater food reward, i.e. symbol 10 (67.9%) rather than symbol 2 (*P* < 0.02; Table 2).

### Quantity discrimination

There was a clear tendency for lovebirds to choose symbols indicating a the greater food quantity over symbols indicating the lesser food quantity. This was the case for all symbol combinations tested except 4 vs. 5. For all combinations with ratios higher than 3, lovebirds chose symbols indicating a greater quantity of food significantly more often than the symbol indicating a lesser quantity of food (Fig. 3a). The proportion of choices for the symbol indicating a greater quantity of food was associated with the ratio of two quantities (model 2: *P* < 0.001). The proportion increased slowly among combinations when ratios were low (between 1 and 3) but increased more substantially between ratios of 3 to 4, with approximately 70% preference for the higher food reward symbol for ratios > 3 (Fig. 3a). That is, a ratio of 3 (i.e., a ratio ≤ 3 vs. > 3) was the general threshold of lovebird discrimination ability (model 3: *P* < 0.01).

**Figure 3.**
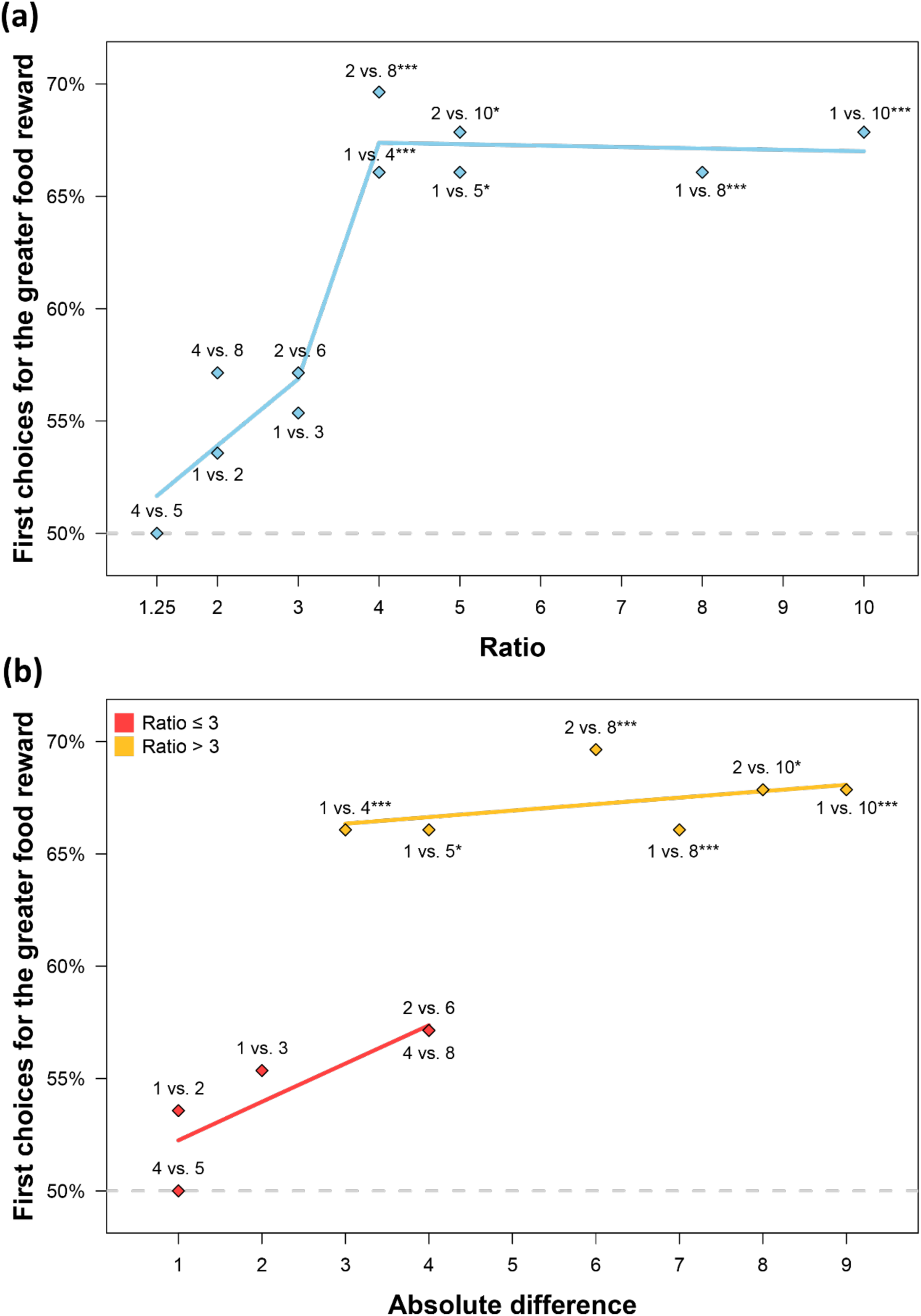
The proportion of individual lovebirds that first opened the cup with the greater food reward as a function of (a) the ratio or (b) the absolute difference indicated between the 2 symbols. The fitted line model in (a): proportion of first choice with the greater food reward ∼ ratio * ratio level. Ratio level: high (> 3) or low (≤ 3). The fitted line model in (b): proportion of first choice with the greater food reward ∼ absolute difference. Significant results in asterisk: * indicates p < 0.05; ** indicates p < 0.01; *** indicates p < 0.001

Absolute differences in food quantities also exhibited an effect on the proportion of instances in which lovebirds chose the symbol indicating the greater quantity of food (model 4: *P* < 0.001). At ratios ≤ 3, choice for the symbol indicating a greater quantity of food increased significantly with absolute differences in food quantity (*P* < 0.001; Fig. 3b). At ratios > 3, this increase was shallower and not significant (*P* = 0.21; Fig. 3b). The influence of absolute differences in food quantities on symbol selection thus also differed between different ratio ranges (ratio ≤ 3 vs. > 3; model 5: *P* < 0.001). When investigating the effect of absolute differences in food quantities within the same ratio band, lovebirds chose the symbol indicating the greater quantity of food more often when presented with combinations that involved larger absolute differences, compared to combinations with smaller absolute differences (Fig. 4).

**Figure 4.**
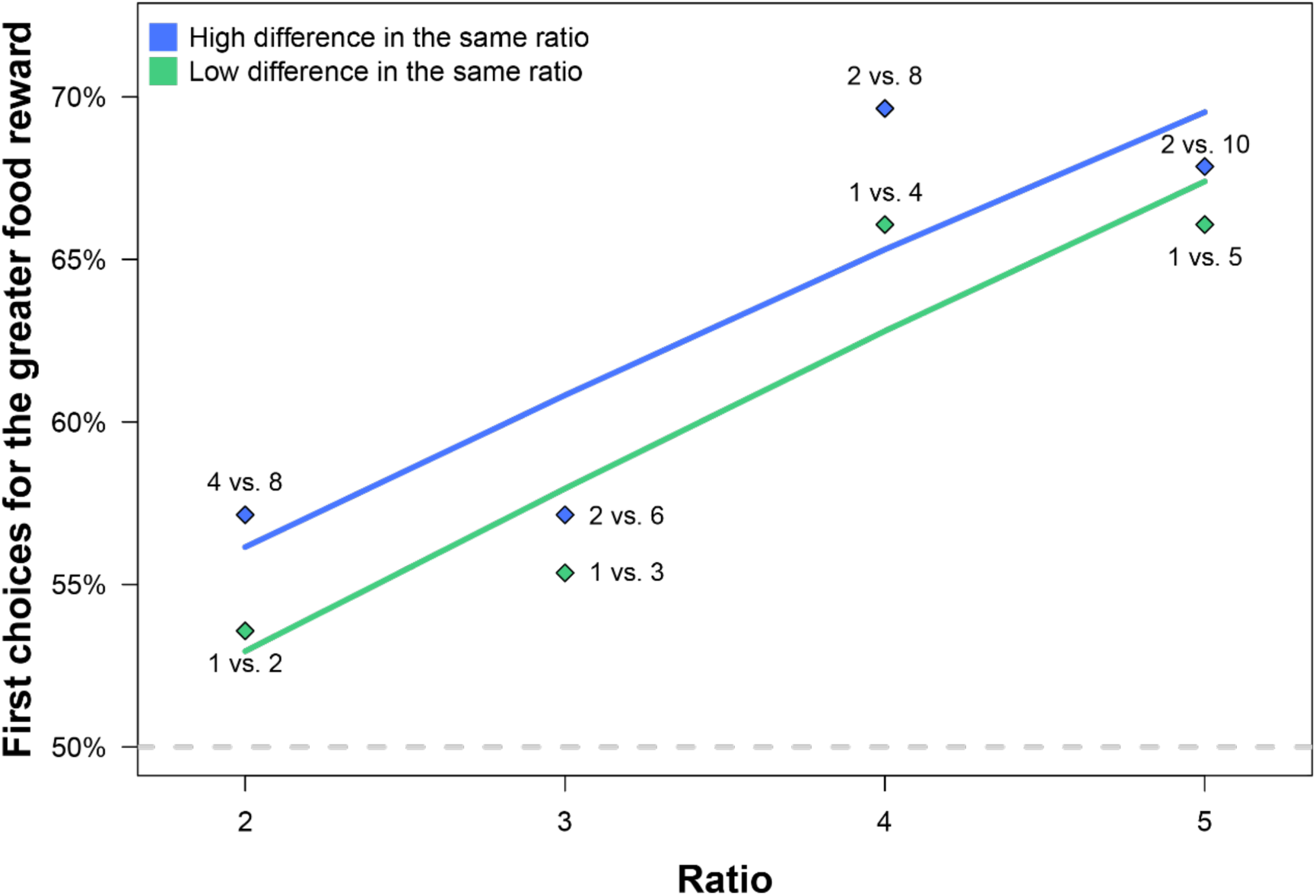
Proportion of first choices for the greater food reward as indicated by symbol combinations of different absolute differences within the same ratio. The fitted line model gives the proportion of first choice with the greater food reward ∼ ratio.

Models detected no significant influence of symbol familiarity (model 6: *P* = 0.629) or of test order (model 7: *P* = 0.104). Across tests of different symbol combinations, the latency of individual birds to choose a feeder was similar (model 8: *P* = 0.301).

### Inter-individual variation in performance

We evaluated the performance of each individual lovebird according to the proportion of first choices it made for the symbol indicating the greater quantity of food across all combinations (Fig. 5). Individuals with the highest tendency to select the greater food reward symbol on their first choice also tended to exhibit the highest tendency to select the greater food reward symbol at low ratio combinations. Sex (*P* = 0.337), age (*P* = 0.597), and body weight (*P* = 0.622) showed no association with an individual’s quantity discrimination performance.

**Figure 5.**
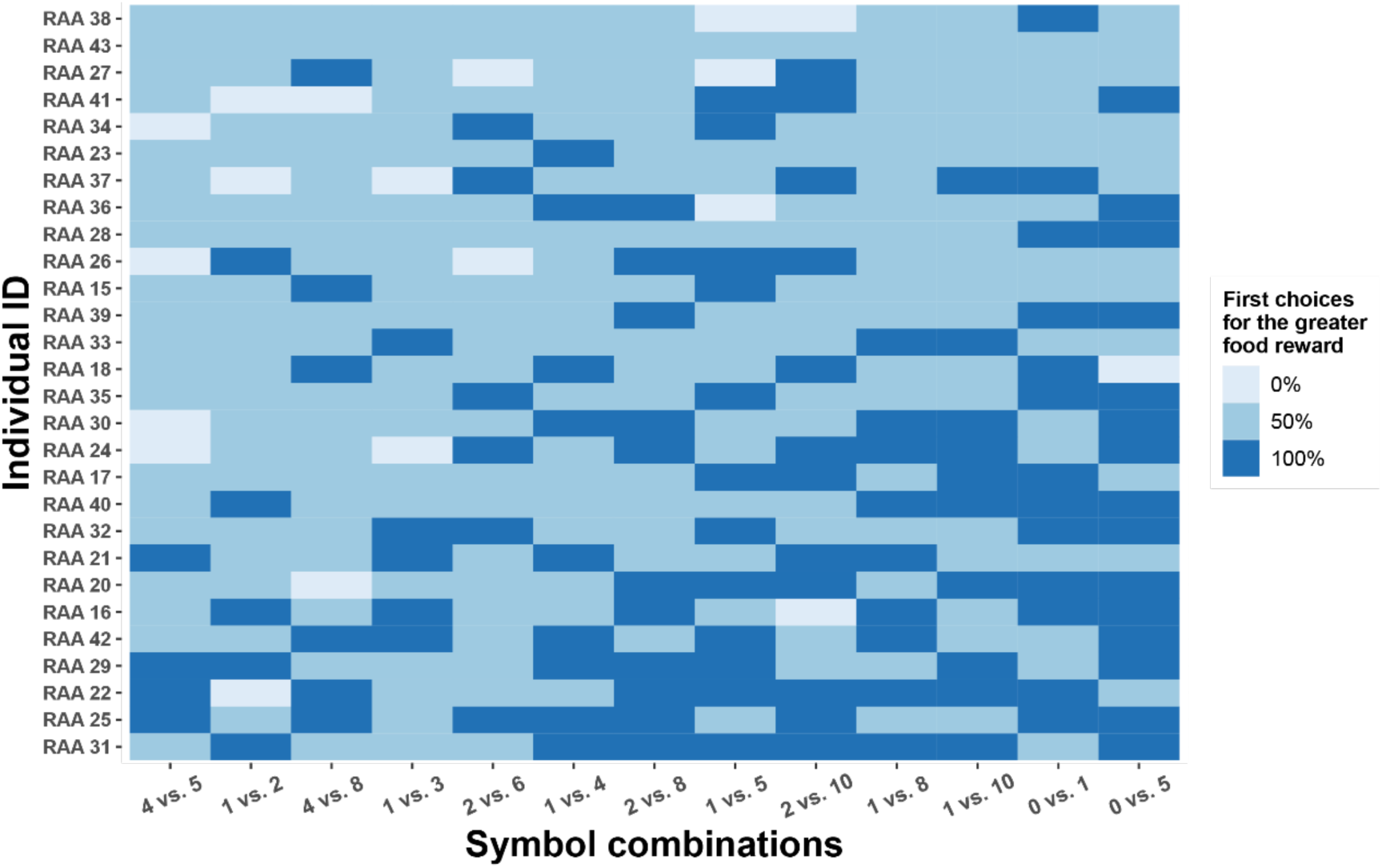
Individual differences in the proportion of first choices for the greater food reward across all symbol combinations. Individuals are ordered according to their overall proportion of first choices for the greater food reward in all combinations, with the individuals with the highest ratio located at the bottom. Symbol combinations are ordered by their ratio, with the lowest ratio displayed on the left.

## Discussion

Our three principal predictions relating to associative learning and inference ability in rosy-faced lovebirds were well supported, with birds proving capable of 1) associating symbols (i.e., object-file symbolism); with 2) “more-less” quantity inference, by deducing food quantities based on their knowledge of this symbol-quantity association; and 3) enhancing their performance in relation to disparity ratio (conforming to Weber’s law) and absolute difference.

### Associative learning of symbol-quantities

Rosy-faced lovebirds were quickly trained to associate a specific food quantity with a ‘|’ tally mark symbol and distinguish this from a ‘O’ symbol, indicating an empty feeder. This rapid associative learning is a trait they share with various other psittaciform species, including red-shouldered macaws (*Diopsittaca nobilis*) and black-headed parrots (*Pionites melanocephala*) [83], as well as species from other avian taxa, including jungle crows [40], and New Zealand robins (*Petroica longipes*) [84]. One highly-trained African grey parrot, named Alex, could even understand the quantities represented by Arabic numbers [85]. In contrast, common pigeons (*Columba livia*) are slow learners, but after 3720 training sessions still proved capable of linking precise numerosities with their corresponding symbols [86]. That some avian species perform better at counting-related associative learning tasks likely reflects inter-species cognitive biases [87]. Furthermore, symbol associations may transcend sensory modalities. For example, Alex the African grey parrot was capable of discriminating a variety of objects based on shape, colour, texture, and auditory labels [88,89].

Individuals of the same species may also differ in performance. While in our study individual differences were unrelated to sex, age or body weight, we did detect that those individuals selecting the greater food reward symbol on their first choice also selected the greater food reward at low ratio combinations. This implies that certain individual lovebirds are more capable of sensitive discrimination than others. This may relate to individual developmental stress trajectories, where Florida scrub-jays (*Aphelocoma coerulescens*) with below average levels of corticosterone as 11-day-old nestlings performed better in associative learning tests as adults, whereas nestlings with above average levels of corticosterone subsequently performed better on reversal learning tests [90]. Similarly, corticosterone exposure during zebra finch (*Taeniopygia guttata*) development improves novel foraging task performance [91]. Individuals can also be influenced by environmental factors, where an unpredictable food supply can cause moderately elevated corticosterone levels in mountain chickadees (*Poecile gambeli*), which correlates with enhanced cache retrieval efficiency and more accurate performance on a spatial memory task [92]. Individual health may also mediate learning ability [93], where general cognitive performance declines with female age in southern pied babblers (*Turdoides bicolor*) [94] and with parasitism [95] in house sparrows (*Passer domesticus*). Fear also affects behaviour and cognitive performance [96]. For example, great tits (*Parus major*) feed sub-maximally in the presence of (model) Eurasian sparrow hawks (*Accipiter nisus*) in order to maintain agility in flight to evade perceived predation [97]. In this regard, lovebirds are gregarious and could have potentially felt exposed when isolated in our trials; nevertheless, we still recorded strong associative learning outcomes.

### More-less quantity inference

Lovebirds were not merely capable of learning to associate symbols with food presence-absence; they could also comprehend the meaning of a food-indicating symbol representing a greater quantity of food, enabling them to infer when the symbol combination on one feeder lid indicated the presence of a greater food reward than that on the other feeder lid. This is an ability lovebirds share with African grey parrots (e.g., Alex; [85]), but also with common pigeons, that can be trained to peck a certain number of times (1, 2, 3 or 4 pecks) on a key that displayed one of several possible numerical symbols [98]. Similarly, jungle crows, can be trained using circle symbols to understand the more-less concept and extrapolate to novel quantity juxtapositions [40].

Rosy-faced lovebirds could further infer and extrapolate the food quantities indicated by novel symbol tallies, exemplifying a capacity for ordinality; an ability shared with rufous hummingbirds (*Selasphorus rufus*) [99]. Lovebirds chose the feeder symbolized with 10 over the feeder symbolized with 2 in equal proportion to 5 vs. 1. This demonstrates a form of transitive inference, where an extrapolation is made based on previous learning, as this related to increasing quantity at the same ratio as a proportion disparity [100]; an ability also observed for trained primates (squirrel monkeys, *Saimiri sciureus* [101], rhesus monkeys [102]). Pepperberg’s [85] work with Alex the grey parrot similarly demonstrated that he could infer the relationship between an Arabic number and a quantity via stimulus equivalence, and understood the ordinal relationship of these numbers.

### Quantity discrimination: more than just ratio

Our results support an analogue magnitude mechanism [103]; that is, as the ratio between feeder pairs increased, the proportion of choices for the feeder containing the greater quantity of food also increased. This demonstrates that lovebirds’ decisions were affected by both the numerical ratio and the stimuli magnitude. This conforms with Weber’s law, in which the discrimination between two quantities is resolved as a function of the ratio between them rather than the absolute difference [69,104,105] according to a Weber-fraction signature (i.e., the ‘just noticeable difference’ between numerosity pairs increased in proportion to numerical magnitudes [106]).

This ability has been widely noted across the animal kingdom [107], from vertebrates to invertebrates [16,69,108–112], involving both non-symbolic [113] and symbolic [114] numerical cognition. Non-symbolic numerical cognition follows at least two distinct systems: ‘parallel individuation’ that encodes the numerical identity of individual items, although this likely applies only to small numbers <4; and the ‘approximate number system’ that encodes the approximate numerical magnitude, or numerosity, of a set, and applies to numbers >4 [115]. Symbols may then acquire semantic meaning through cognitive mapping onto the approximate system for the non-symbolic representation of number (Approximate Number System or ANS; [116]). The analogue magnitude system enables individuals to make rapid comparisons without quantity limits [117] and enables both continuous and discrete amounts to be compared [69]. In our experiment, lovebirds could not directly observe quantities or volumes and form an object-file representation on that basis, rather they learned to associate a tally mark with a specific food quantity as a one-to-one correspondence between object files [38]

In addition to analogue magnitude effects, we observed that an absolute difference between feeder pairs promoted the rate at which lovebirds chose the more rewarding feeder (Figure 4), which has also been observed in African grey parrots (19). This is an effect many studies overlook or judge inconsequential ([44]; but see [118,119]), although Neider [44] has demonstrated that not every vertebrate species he tested for numerical cognition was able to flexibly discriminate absolute numerosity, suggesting that qualitative differences in numerical intelligence occur between vertebrates.

Furthermore, our modelling revealed a significant interaction term between feeder quantity ratio and absolute difference, such that the impact of absolute differences diminishes at higher ratios, which suggests that the role of absolute quantity is secondary. With large ratio disparities the magnitude of difference between quantities likely overwhelms a perception of absolute difference, and the absolute value of either option is perceived as sufficiently plentiful [120]. As ratio diminishes, it becomes increasingly challenging to distinguish between two quantities, so the absolute difference can assist in quantity discrimination. In support of this we found a notable divergence at different ratio intervals (i.e., ratio ≤ 3 vs. ratio > 3), evident as a statistically significant ability for lovebirds to distinguish different quantity combinations, where ratios and absolute differences influenced the proportion of instances where individuals chose the feeder containing the greater quantity of food (Fig. 3). This suggests that, across individuals, lovebirds have an average discrimination threshold around a ratio limit (ratio 3); however, this varied between individuals, with those individuals scoring a more rewarding overall performance apparently being either better able to differentiate quantities of lower ratios or exhibiting a higher motivation to choose optimally. Such inter-individual variation in cognitive capacity can be critical for survival and reproductive success [121–123]. This threshold varies among species. In African grey parrots, Aïn et al. [69] reported all three of their subjects successfully discriminated between visually available food rewards at a ratio 1.5 (2 vs. 3) and higher (up to 1 vs. 5). For domestic cats (*Felis catus*), this threshold is greater or equal to 2.5 (2 vs. 5) [124], and greater or equal to 1.2 (5 vs. 6) for jungle crows [40].

### Conclusion

The transcendent view across cognitive neuroscience, child psychology, and animal cognition is that there is a biological capacity specifically evolved for understanding numerical quantities and arithmetic, where quantical cognition (quantity discrimination) provides biologically evolved preconditions for numerical (exact, symbolic) cognition [125]. As with other bird species, most notably Corvidae and Psittacidae, we found that rosy-faced lovebirds were capable of advanced quantity assessment and numeracy, where avian capacity for associative learning is a function of the telencephalic area nidopallium caudolaterale (NCL), analogous to the independently evolved prefrontal cortex in mammals [126–128].

## Ethical notes

All procedures were approved by the Committee on the Use of Live Animals in Teaching and Research (CULATR; approval number: 5883-21), and under a Department of Health Animal (Control of Experiments) Ordinance Chapter 340 permit ((21-1146) in DH/HT&A/8/2/3 Pt.32).

## Data accessibility

Data and code are available online as supplementary materials.

## Competing interests

The authors declare no conflicts of interest.

## Author Contributions

Conceptualization, S.Y.W.S.; Methodology, S.W., E.S.K.P., S.Y.W.S.; Investigation, S.W., K.C.H.L., V.W.T.S., C.Y.C.H.; Formal analysis, S.W.; Visualization, S.W.; Writing – original draft, S.W.; Writing – review & editing, S.W., C.N., C.D.B., S.Y.W.S., and all other authors; Funding acquisition, S.Y.W.S.; Resources, E.S.K.P., S.Y.W.S.; Project administration, S.Y.W.S.; Supervision, S.Y.W.S.

## Acknowledgements

We would like to express our gratitude to the CCMR staff Ellen Sai Nam Lo, Mei Ying Wu, and Chun Kiu Lo for taking care of our lovebirds. We thank Wynne Wan Hei Ting, Sarah Hiu Wa Kwok, Dino Chun Yin Tsui, and Joyce Yuk Shan Lam for assistance during the experiments.

## Notes

### Competing Interest Statement

The authors have declared no competing interest.

